# Molecularly defined olfactory bulb neuron subtypes form broad and selective pathways to cortex

**DOI:** 10.64898/2026.10.27.762102

**Authors:** Sourjya Baibhabee Nath, Patrick Niall Stoney, Maria Inês Ribeiro, Shin-Yu Lee, Izumi Fukunaga

## Abstract

Early sensory processing decomposes signals into elemental features conveyed through parallel pathways defined by cell type. In the olfactory bulb, molecular profiling identified subtypes of mitral and tufted cells, but their morphology and projection patterns remain unclear. Using intersectional Cre targeting, retrograde labelling, and RiboTag sequencing, we linked subtype morphology and projections to marker expression. The same cortical area recruited morphologically distinct neurons depending on the Cre line. M1 neurons, enriched in the accessory olfactory bulb, were densely labelled from cortical amygdala; M2 neurons, with deep somata and dendrites, projected widely but were selectively recruited from posterior piriform cortex. T1 cells projected to the olfactory tubercle cap, and superficial ET2 cells to the anterior olfactory nucleus pars externa. Deep, large neurons coupled weakly to sniffing, unlike most extracellularly recorded units, suggesting these recordings under-sample them. Bulb output subtypes thus form parallel channels routing information to broad and selective cortical targets.

## Introduction

Sensory systems transform high-dimensional, complex inputs into distinct features that can be represented and transmitted by specialized neural pathways (Roska & Meister, 2014; Sanes & Masland, 2015; Wassle, 2004). Parallel processing is a central organizing principle of this transformation, enabling multiple stimulus features to be extracted and conveyed simultaneously (Diamond *et al*, 2008; Grothe *et al*, 2010; Nassi & Callaway, 2009; Sanes & Masland, 2015; Wässle, 2004). The olfactory bulb is the first central relay for odor information, where sensory input from the nasal epithelium is transformed into outputs that are then distributed across multiple higher brain regions. However, key questions remain: How is olfactory information routed to distinct downstream pathways, and how do they correspond to molecularly defined olfactory bulb output neuron subtypes? Resolving these questions requires determining how genetic identity relates to cellular morphology, projection organization, and physiology.

Previous studies identified two broad classes of olfactory bulb output neurons, mitral and tufted cells, which differ in morphology, biophysiological properties, local circuit interactions, long-range projections, and neuromodulation (Chae *et al*, 2022; Chen *et al*, 2022; Eiting & Wachowiak, 2020; Fukunaga *et al*, 2012; Fukunaga *et al*, 2014; Haberly & Price, 1977; Igarashi *et al*, 2012; Kapoor *et al*, 2016; Lindeman *et al*, 2024; Luskin & Price, 1982; Nagayama *et al*, 2014; Otazu *et al*, 2015; Scott, 1981; Yamada *et al*, 2017). However, morphological and molecular evidence points to a finer distinction. For example, classical single-cell morphological reconstructions show that the neurons differ not just in the soma depth, but also where the lateral dendrites stratify in the fiber-rich external plexiform layer (EPL), occupying seemingly discrete deep, middle, and superficial portions in a manner reminiscent of the stratification of the dendritic arbors of retinal ganglion cells (Macrides & Schneider, 1982; Mori *et al*, 1983; Orona *et al*, 1984). Further single-nucleus RNA sequencing combined with retrograde tracing revealed 3 subtypes of mitral cells (M1-3), and 5 types of tufted cells (T1, ET1-4) based on gene expression profiles (Zeppilli *et al*, 2021). It is not clear if and how these molecularly defined subtypes relate to morphological classes and long-range projection patterns, and if they differ electrophysiologically.

Here, we used an intersectional approach combining conditional retrograde labeling from six downstream targets in two Cre-driver lines. With morphological analysis and RiboTag-based molecular profiling of olfactory bulb projection neurons, we identify a correspondence between molecularly defined projection neuron subtypes and their downstream projection targets. Molecularly defined M1 neurons are preferentially labeled by retrograde tracing from the cortical amygdala and sparsely from the anterior olfactory cortex, whereas M2 neurons are labeled from a broader set of olfactory cortical regions. T1 neurons differ from ET types in the depth of their lateral dendrites and in projections to the olfactory tubercle. Neurophysiology with opto-tagging revealed that M2 neurons show larger spike amplitudes and weaker sniff coupling. Together, these results relate molecular identity to cellular morphology and long-range projection patterns and suggest that they contribute physiologically distinct channels of olfactory processing.

## Results

### Tbx21-Cre and Cck-Cre provide genetic access to olfactory bulb output neuron subtypes

We wished to find a method to label and study the neurophysiology of subtypes of mitral and tufted cells. Since these subtypes may target different downstream regions, we first tested if retrograde labeling from different olfactory cortices could be used to selectively label mitral and tufted cell types. We injected retrogradely transported adeno-associated virus (AAVrg) into the anterior and posterior parts of the olfactory cortex for unconditional expression of eYFP for morphological analysis. The targets were the anterior olfactory nucleus (AON), olfactory tubercle (OT), and piriform cortex (PIR). Two weeks after AAVrg injection, many eYFP-positive neurons were observed in the olfactory bulb. However, many labelled cells did not resemble mitral or tufted cells, with small somata and thin dendrites with puncta, especially for the AON and OT injections (Fig 1A,B).

**Figure 1:**
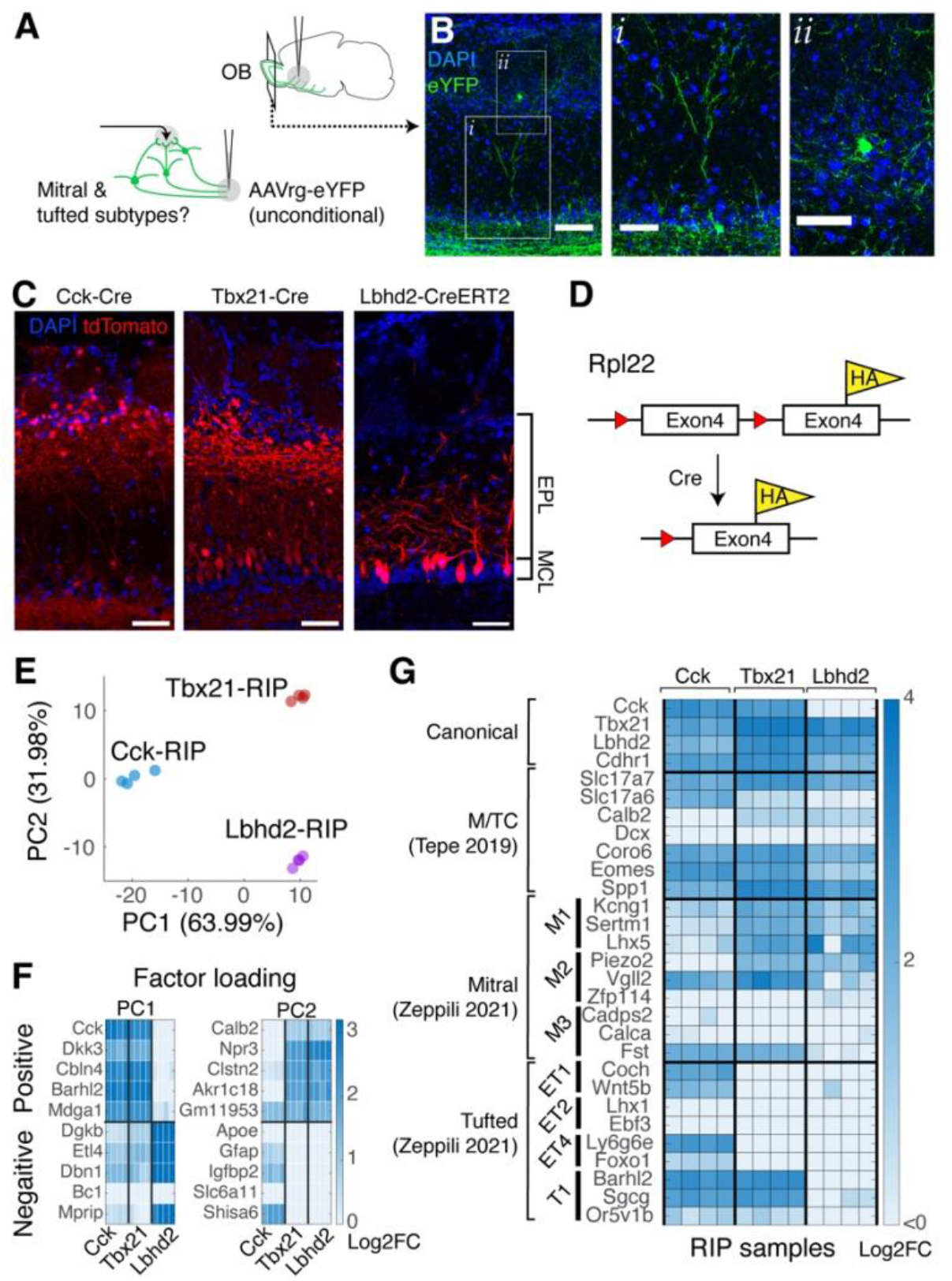
Molecular constraints to study target-specific olfactory bulb output neurons. **A**. Left, experimental configuration, showing injections of unconditional AAVrg for eYFP expression in olfactory cortex. **B**, Examples of fluorescently labeled neurons with unconditional rAAV. Images (i) and (ii) are from the area demarcated in the left image. Scale bar = 0.5 mm. **C**, Examples of tdTomato labeling pattern in OB layers in Cck-Cre::Ai14, Tbx21-Cre::Ai14, and Lbhd2-CreERT2::Ai14 mice. EPL = external plexiform layer, MCL = mitral cell layer **D**, RiboTag construct, adapted from (Sanz et al, 2009). In Cre-positive cells, the endogenous exon 4 of Rpl22 is excised and a modified exon 4 encoding a 3xHA-tag is expressed. **E**, Plot of the first two principal components of RNA-seq data from Cck-RIP, Tbx21-RIP, and Lbhd2-RIP samples. **F**, Factor loadings for the PCA analysis in **E**. Log2 fold change in expression (RIP vs. input) for the top 5 genes in the positive and negative factor loadings for the first two principal components are shown. **G**, Enrichment of published markers of mitral and tufted cells and their subtypes (Tepe *et al*, 2018; Zeppilli *et al*., 2021). Plotted values are log2FC enrichment in RIP samples relative to input.

Since unconditional expression may label immature, GABAergic neurons migrating towards the olfactory bulb through the rostral migratory stream (Lledo *et al*, 2006; Lois & Alvarez-Buylla, 1994), we turned to conditional labeling using three existing Cre-driver lines. Cck-Cre (Taniguchi *et al*, 2011), Tbx21-Cre (Haddad *et al*, 2013), and Lbhd2-CreERT2 (Koldaeva *et al*, 2021) are commonly used to study OB output neurons. As previously reported, when the Cre lines were crossed with Ai14 tdTomato reporter mice, red fluorescence was restricted to mitral and tufted cells, with bias towards the more superficial, tufted cells in Cck-Cre, whereas both deep and superficial neurons (mitral cells and tufted cells) were labelled in Tbx21-Cre (Fig 1C). Inducible recombination with Lbhd2-CreERT2 resulted in sparse but morphologically selective labeling of mitral cells. This morphological analysis indicates that the three Cre-driver lines constrain the labeling to the principal neurons, with overlap in the labelled cell types.

To further validate the Cre lines, transcriptomic analysis was carried out to test if the Cre-expressing cells in Cck-Cre, Tbx21-Cre, and Lbhd2-CreERT2 lines span the known subtypes of mitral and tufted cells. We turned to bulk RNA-seq using the Ribotag mouse, which carries a modified *Rpl22* allele (Sanz *et al*., 2009). In this design, the Cre-recombinase excises the endogenous exon4 of *Rpl22*, leading to the expression of a modified exon4 that contains a 3xHA tag (Fig 1D), allowing cell type-specific immunoprecipitation of ribosomes together with their associated mRNAs (RNA-immunoprecipitation, RIP). Immunoprecipitated, ribosome-bound RNAs were sequenced and compared to input samples. Non-RPL22-HA-expressing control samples were used to exclude non-specific RNA enrichment.

mRNA enrichment patterns were distinct for Cck-, Tbx21- and Lbhd2-RIP samples, as demonstrated in the clear separation on the principal component axes (Fig 1E,F). RIP samples were also well-separated from both the input and control IP samples (Supplementary Fig.1). Transcripts previously identified as markers for the subtypes of mitral and tufted cells (Zeppilli *et al*., 2021) were enriched in the RIP samples, and overlapped between the three driver lines (Fig 1G). Fewer transcripts were enriched in Lbhd2-RIP samples, most of which were also enriched in Tbx21-RIP samples (298/369 genes, or 80.8%; Supplementary Fig. 1B), consistent with a morphologically more selective population (Fig. 1C; (Koldaeva *et al*., 2021)). Overall, these results demonstrate that Cck- and Tbx21-positive cells together cover the mitral and tufted cells comprehensively. But because of the overlap in the enrichment patterns of subtype-specific transcripts, an additional method such as projection mapping is needed to selectively analyze heterogeneous olfactory bulb output neurons.

### Intersectional targeting reveals diverse laminar morphological profiles

To analyze the distribution of mitral and tufted cell subtypes in Cck- and Tbx21-positive neurons, we next investigated the projection patterns of OB outputs. Retrograde AAV encoding Cre-activated RPL22-HA and eYFP (AAVrg-FLEXon-EF1A-RPL22-3xHA-IRES-EYFP-WPRE) was injected into 6 locations spanning the anterior-posterior extent of the olfactory cortex (Fig. 2A, B). AAVrg was injected into the external and the ventrolateral parts of the anterior olfactory nucleus (AONpe and AONvl, respectively), the cap region of the olfactory tubercle (OTcap), anterior piriform cortex (aPIR), posterior piriform cortex (pPIR), and the cortical amygdala (CoA) of Cck-Cre or Tbx21-Cre mice (Supplementary Fig. 2). Two weeks after AAVrg injection, eYFP labeling in the olfactory bulb was assessed (Fig. 2C,D). The labelled neurons showed the typical morphology of mitral and tufted cells, with medium to large somata (interquartile range of soma areas = 77.73 – 166.24 μm^2^ for Cck-Cre and 106.4 – 227.74 μm^2^ for Tbx21-Cre), thick apical dendrites and with smooth lateral dendrites in the external plexiform layer. As expected, injections into the anterior regions labelled more superficial cells, in Cck-Cre mice, particularly in AONpe (Supplementary Fig. 3; percentage of labelled somata in superficial EPL = 59.2 ± 10.2%; p = 4.33 x 10-43, F = 92.35, d.f. = 11; 1-way ANOVA; n = 11 injections for AONpe). Injections caudal to AON labelled few superficial cells (percentage of labelled somata in superficial EPL = 6.5 ± 7.1% and 2.8 ± 2.9% for Cck-Cre and Tbx21-Cre, respectively). All injections in Tbx21-Cre mice densely labelled deep cells (proportion of labelled somata in deep EPL = 51.87 ± 4.41 % for AON and 74.50 ± 7.23% for caudal regions). AAVrg injections caudal to AON in Tbx21-Cre mice consistently labelled both somata and lateral dendrites in deep EPL. In Cck-Cre mice, labeling following caudal injections comprised deep cells (proportion of labelled somata in deep EPL = 68.01 ± 8.76% for caudal regions) but only sparse labeling was observed (Supplementary Fig. 3). CoA injections resulted in dense labeling of the accessory olfactory bulb (Supplementary Fig. 4). OTcap injections in Cck-Cre mice consistently labelled deep cells with dendrites in the middle portion of EPL (Fig. 2C,D).

**Figure 2:**
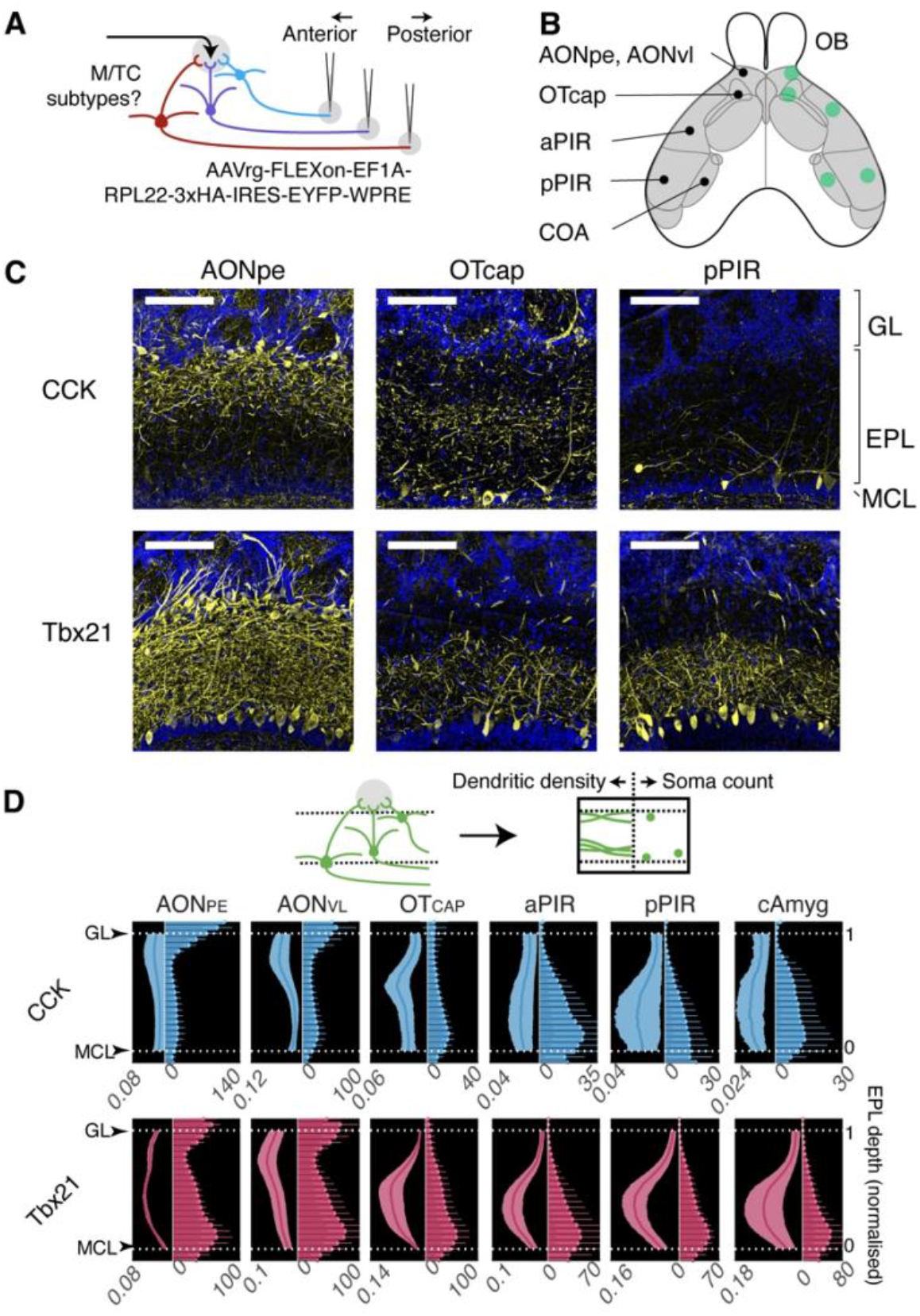
Morphologically heterogeneous OB output labeling with molecularly constrained projection mapping. **A**, Experimental configuration. Retrogradely transported AAV encoding Cre-activated RPL22-HA and eYFP was injected into regions spanning the anterior to posterior extent of the olfactory cortex. Cre-drivers used were Cck-Cre and Tbx21-Cre. **B**, Schematic representations of injection targets. AONpe= anterior olfactory nucleus pars externa, AONvl = ventral lateral AON, OTcap = cap region of the olfactory tubercle, aPIR = anterior piriform, pPIR = posterior piriform cortex, CoA = cortical amygdala. **C**, Confocal images showing the labelled fluorescence patterns in the olfactory bulb, following injections into AONpe, OTcap, and pPIR in Cck-Cre and Tbx21-Cre mice. Scale bar = 0.1 mm. GL = glomerular layer, EPL = external plexiform layer, MCL = mitral cell layer. **D**, Quantification of labelled soma counts (leftward axis) and dendritic eYFP signal density (rightward aixs) relative to the boundaries of the EPL. Mean and SD are shown.

### Intersectional RiboTag profiling reveals overlapping but distinct subtype-marker enrichment patterns

The morphological diversity above suggests that a combination of target-specific retrograde labeling and Cre-driver lines can distinguish subtypes of mitral and tufted cells. An important question that follows is whether and how the subtypes that were defined previously by gene expression patterns (Zeppilli *et al*., 2021) correspond to our data. To address this, we injected AAVrg-FLEX-RPL22HA-eYFP into 6 regions of either Cck-Cre or Tbx21-Cre mice for RNA-IP and RNA-seq analysis (Fig. 3A). The injection conditions were Cck-AONpe, Cck-AONvl, Cck-OTcap, Cck-CoA, Tbx21-OTcap, and Tbx21-pPIR, to cover a range of superficial to deep mitral and tufted cells. After sequencing, we compared the RIP enrichment of marker transcripts in the different injection conditions (Fig. 3B). Transcripts commonly expressed by immature interneurons, GABAergic neurons, or glial cells were depleted in these samples (Supplementary Fig. 5; Table 1).

**Figure 3:**
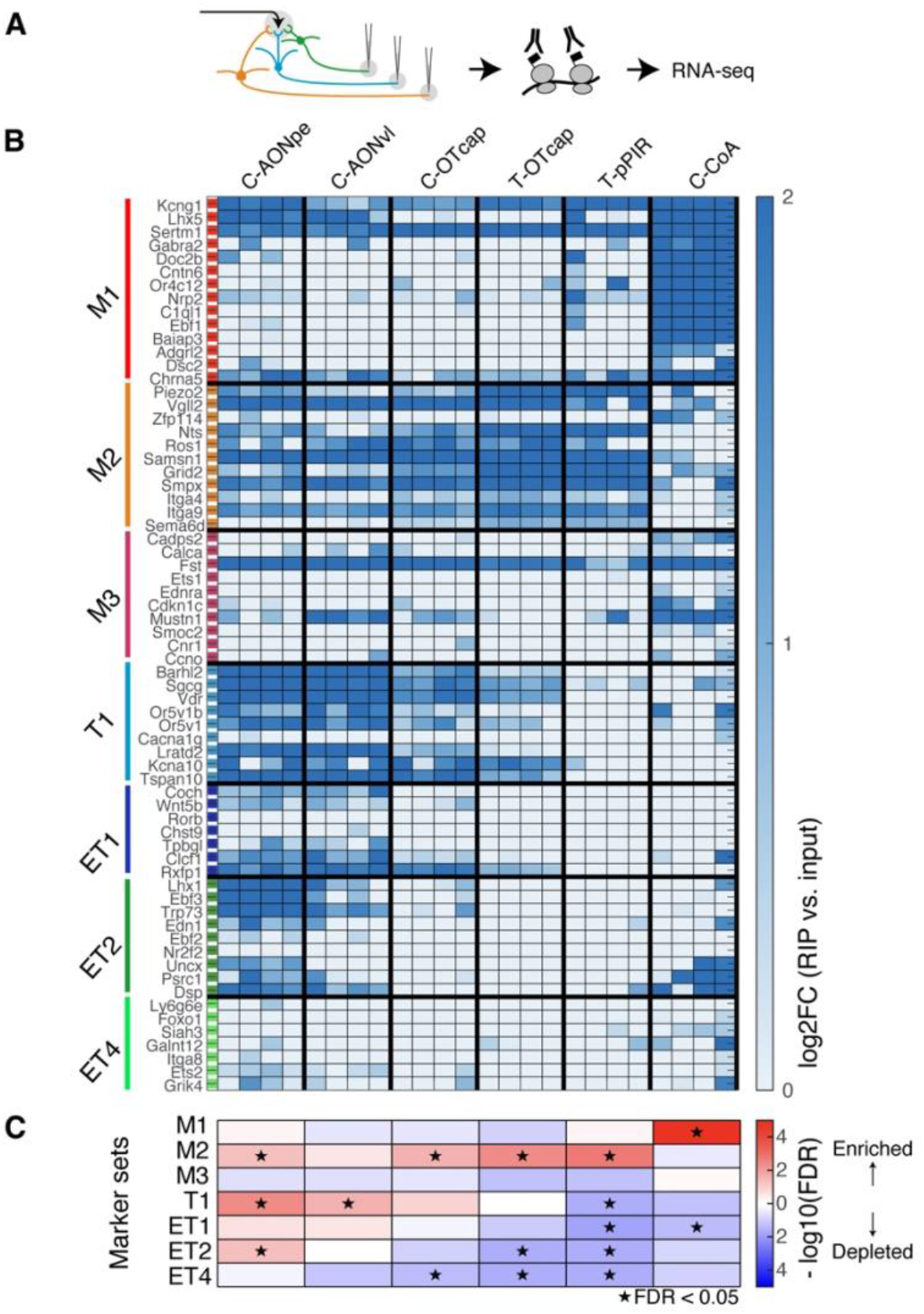
Correspondence of projection targets with molecularly defined MTC subtypes. **A**, Experimental configuration. Conditional expression of HA-tagged RPL22 using AAVrg. Immunoprecipitated mRNA was sequenced. **B**, Heatmap of log2FC enrichment in RIP samples vs. input of subtype-specific marker genes (Zeppilli et al., 2021). **C**, Correlation Adjusted MEan RAnk gene set test analysis (Wu & Smyth, 2012) of the subtype-specific markers. Heatmap of -log10 of false discovery rate (FDR; Benjamini-Hochberg), with enrichment and depletion in red and blue hues, respectively.

**Table 1:**
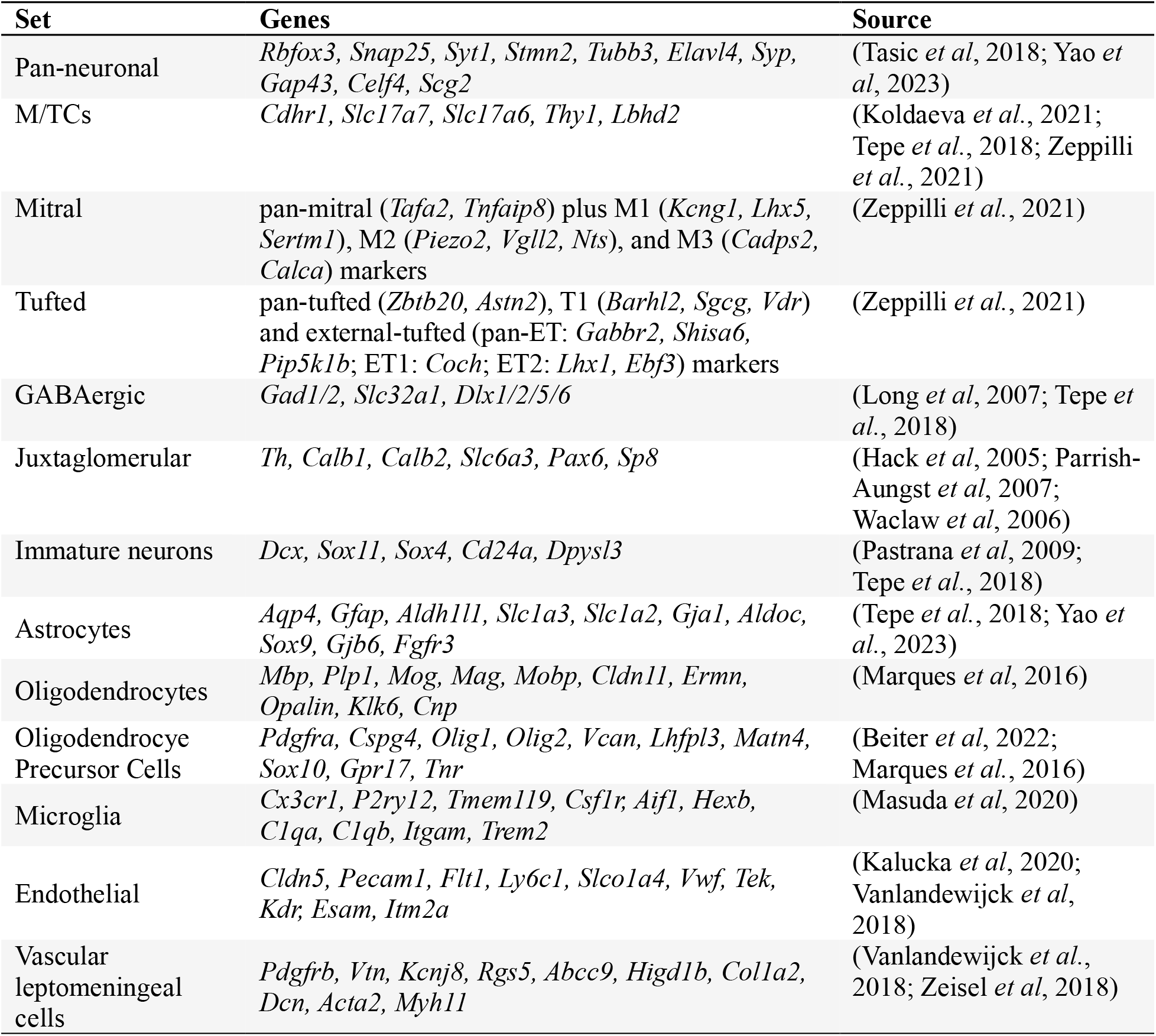
Gene sets used to assess cell type-specific transcript enrichment reported in Supplementary Figure 5.

We found that M1 marker transcripts were enriched most consistently in CoA-projecting neurons (Fig. 3B,C; Table 2; FDR = 1.92 x 10^-16^, 14 transcripts), with enrichment of a few markers in Cck-AONpe samples. Analysis of publicly available *in situ* hybridization data from the Allen Brain Atlas revealed that many transcripts that were enriched in Cck-CoA samples (Table 3) are consistently expressed in AOB (Supplementary Fig. 6). M2 markers were enriched in most injection conditions (Fig. 3B,C; FDR for Cck-AONpe, Cck-AONvl, Cck-OTcap, Tbx-OTcap, Tbx-pPIR = 0.044, 0.27, 0.023, 0.0039, 0.0015, respectively; 11 transcripts) but tended to be depleted in Cck-CoA (FDR = 0.33). For pPIR and OTcap injections in Tbx21-Cre mice, M2 markers were the only set that were enriched in RIP samples (Fig. 3B,C). ET2 markers were most abundant in AONpe-projecting Cck neurons (FDR = 0.045, 9 transcripts) and distinguished by depletion in the Tbx21-OTcap and Tbx21-pPIR samples (Fig. 3B,C; FDR = 0.022 and 0.023, respectively). T1 markers were enriched in samples following the anterior, AON injections (FDR for Cck-AONpe and Cck-AONvl =0.0039 and 0.022, respectively; 9 transcripts), and also tended to be enriched in OTcap samples. Altogether, projection-specific conditional expression of RPL22-HA with eYFP labeling allowed the anatomical identification of mitral and tufted cell subtypes.

### Intersectional optotagging reveals differences in spike amplitude and sniff locking

Do the morphological differences correspond to physiological differences? To address this question, we turned to extracellular electrophysiology combined with opto-tagging. Retrograde AAV conditionally expressing ChR2-eYFP was injected into AONpe or OTcap of Cck-Cre mice, or aPIR of Tbx21-Cre mice (Fig. 4A,B). Based on our morphological and transcriptomic analyses, these injections cover a gradient, starting with a mixed population dominated by superficial, tufted cells towards deep, mitral cells. Extracellular spikes were recorded using a 32-channel optrode from the olfactory bulb of anesthetized mice that breathed naturally (2.49 +/- 041 Hz, n = 15 mice). To evoke activities in ChR2-expressing neurons, brief, 5-ms pulses of blue light were presented every 5 seconds. Some units generated consistent, short-latency spikes in response to the light pulses (Fig. 4C). These are referred to as opto-tagged units.

**Figure 4:**
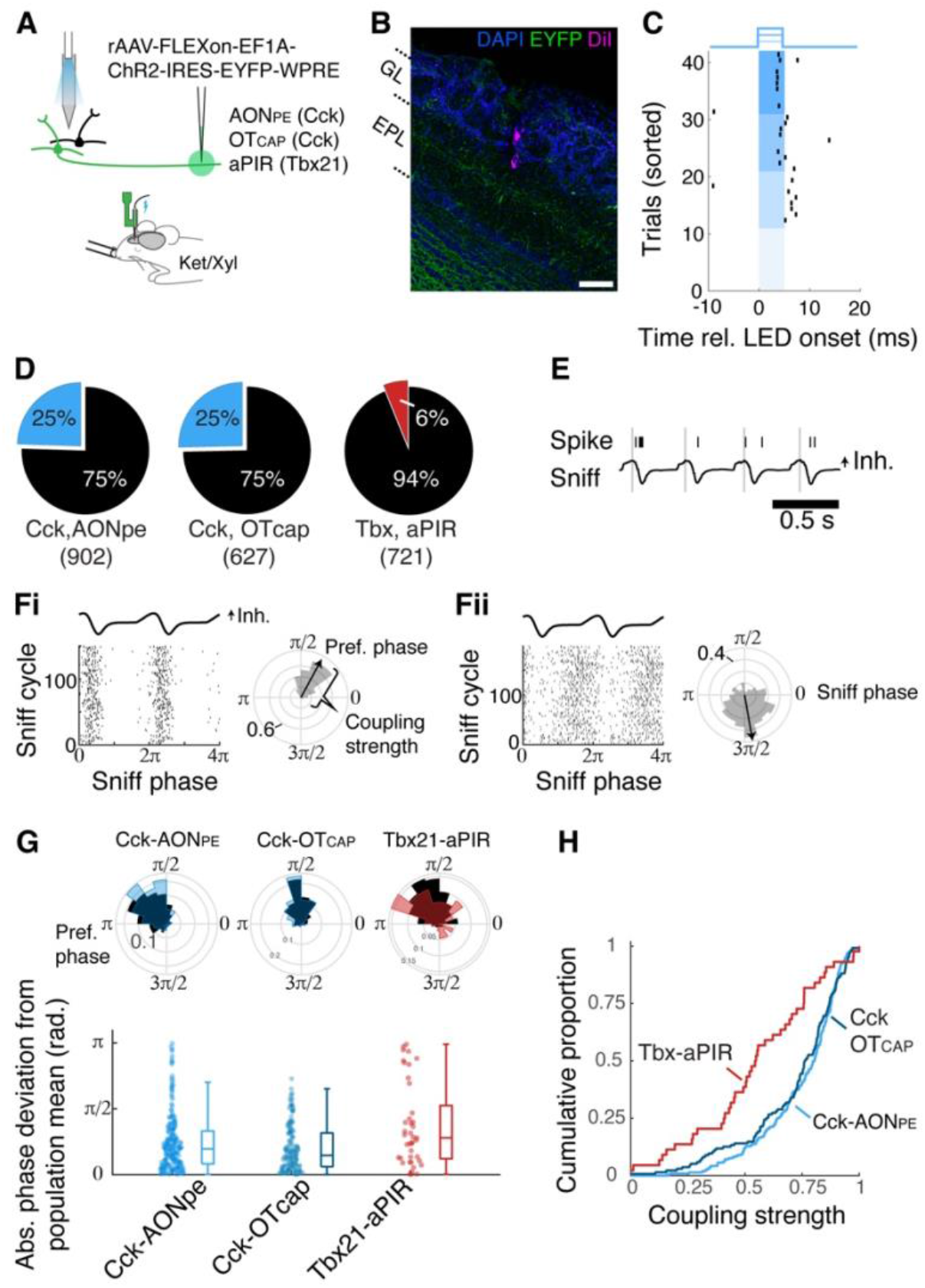
Sniff-locking of piriform-projecting Tbx21 neurons deviates from the population average. **A**, Experimental configuration. rAAV for conditional expression of ChR2-IRES-EYFP injection was targeted to either AONpe or OTcap of Cck-Cre mice, or aPIR of Tbx21-Cre mice. Acute extracellular electrophysiology was conducted from the olfactory bulb under ketamine/xylazine anaesthesia with a 32-channel probe with a 50/62.5 mm fiber optic cannula 0.1 mm above the highest recording site. **B**, Example of post-hoc histology showing DiI labeling the electrode track, eFYP from labelled neurons, and DAPI delineating the OB layers. Scale bar = 0.1 mm. **C**, Spike raster plot for an example opto-tagged neuron around the time of light presentation, sorted by the irradiation level used (blue shaded areas indicate the time of light presentation; from low to high, 0, 0.012, 0.22, and 0.34 mW). **D**, Percentages of opto-tagged units (colored segment) for injection condition. The numbers in brackets indicate the total numbers of units. n = 4, 6, 5 mice for the 3 experiments, respectively from left. **E**, Example nasal flow ("sniff", lower trace) and spike times for an example unit (black ticks). A sniff cycle is defined from one inhalation peak to next (gray vertical lines). Air flow in the positive direction corresponds to inhalation ("Inh.") **F**, Left, Spike raster aligned to the sniff cycle. The timings of spikes are expressed as the phase within each cycle, in radians. The phase of spikes relative to the sniff cycle is summarized as polar histograms, and the resultant vector is shown as black arrows. The coupling strength is the length of this resultant vector. Bi and Bii are different examples. **G**, Top, Summaries of preferred phase, plotted as polar histograms, for all recorded units (black) and opto-tagged units (colored segments). Bottom, phase deviations for the opto-tagged units from the mean phase from all units from the same animal. **H**, Cumulative histograms of sniff coupling strengths for the opto-tagged units for the indicated injection condition.

In Cck-Cre mice, approximately 25% of the sorted units were categorized as opto-tagged for both AONpe and OTcap injections (Fig. 4D; 24.5% or 221/902 units from 4 mice and 25.4% or 159/627 units from 6 mice, respectively), while only 6% of units were opto-tagged in Tbx21-aPIR condition (41/721 units from 5 mice). The low level of recruitment for the Tbx21-aPIR condition is not explained by the difference in labeling densities or the depths recorded. We then asked if the opto-tagged units are physiologically different.

We then wished to determine if there are physiological differences related to local connectivity. We analysed the sniff-coupled activity, as its timing can be sculped by local inhibitory circuits (Fukunaga *et al*., 2014). Since the cell type-dependent difference in sniff-coupling is most striking in the absence of odors, and the phase relationship is sensitive to changes in the sniff frequency, spiking activity from the baseline period from anesthetized mice were analysed (Fig. 4E-H). To assess general coupling, we measured the preferred phase and coupling strength for each unit (the direction and amplitude of the resultant vector of spike histogram; Fig. 4F).

Interestingly, the preferred phase of the opto-tagged units in both Cck-AONpe and Cck-OTcap injections largely aligned with the population (Fig. 4G,H; median deviations = 0.62 with interquartile range [0.26 - 1.04] radians and 0.46 [0.19 - 0.99] radians, respectively; n = 221 and 159 units for Cck-AONpe and Cck-OTcap), while the preferred phase of opto-tagged units in Tbx21-aPIR injection deviated significantly from the population mean (0.88 [0.38 - 1.65] radians; n = 41 units; p = 0.0057; F = 5.23, d.f. = 2, nested ANOVA). Further, the opto-tagged units in Tbx21-aPIR injection showed much weaker locking than those in Cck-AONpe and Cck-OTcap cases or in the general population (p = 1.75 x 10-5 and 3.75 x 10-4, relative to Cck-AONpe and Cck-OTcap, respectively; Kolmogorov-Smirnov test, D = 0.40 and 0.35, respectively). The results indicate that tufted cells show robust sniff coupling, the timing of which aligns with that of most extracellularly sampled units, while deep mitral cells deviate from the majority, generating spikes with a slightly shifted timing and weaker coupling.

## Discussion

Determining whether olfactory processing is implemented in fine-grained parallel pathways required a direct comparison of molecular profiles, cell morphology, and long-range projection patterns. Our data revealed that the molecularly defined mitral and tufted subtypes differ in their morphology, including the stratification of their lateral dendrites within the olfactory bulb layers, and have overlapping but distinct cortical projections (Figure 5). The results imply that the target alone is not the only determinant and that the genetic identity predicts a coordinated anatomical phenotype. In addition, the implication of this architecture is that the results of subtype-specific local computations in the olfactory bulb may be combined differently by distinct downstream regions.

**Figure 5:**
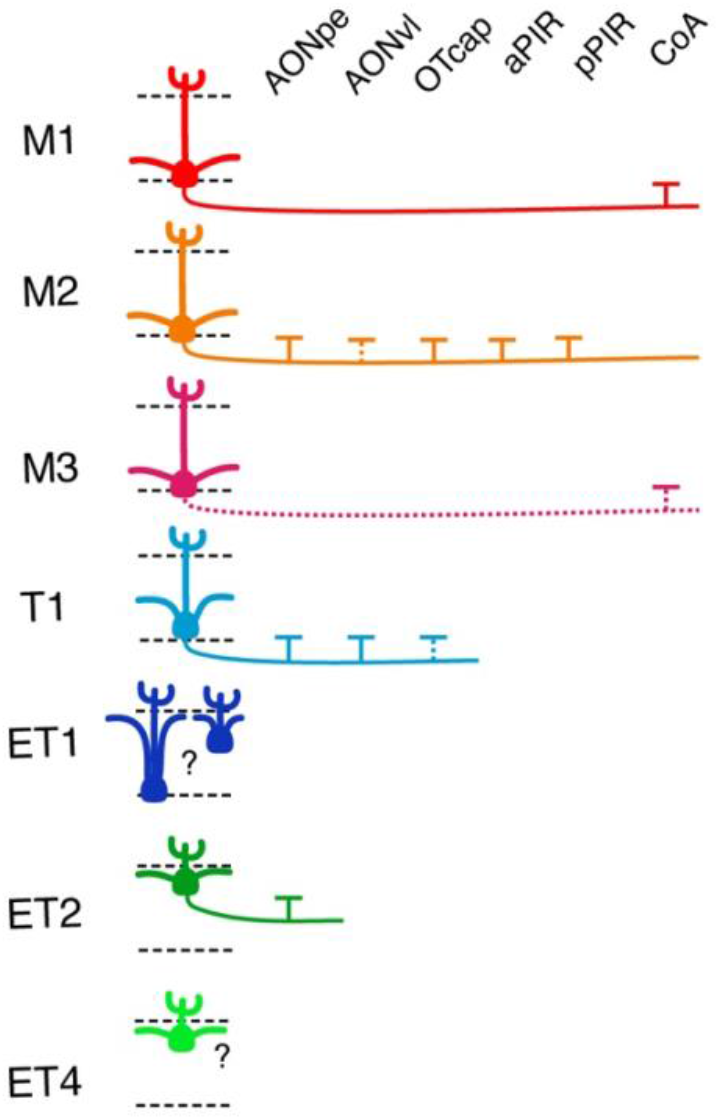
Graphical summary of the findings. Molecularly and morphologically distinct subtypes of olfactory bulb output neurons target a distinct set of downstream targets, suggesting a unique integration of olfactory information in secondary steps of olfactory processing.

Given that a consensus in the field is paramount, we provide direct comparisons of our results to previously reported subtypes, based on comprehensive molecular profiling. Previous studies reported M1 neurons to be recruited from anterior locations (Zeppilli *et al*., 2021). While some genetic markers of M1 neurons were enriched in cells projecting to anterior cortical areas, they were most consistently enriched in CoA-targeting populations. According to Allen Brain Atlas ISH data, M1 markers are particularly enriched in the accessory olfactory bulb. M2 markers were enriched in cells projecting to most regions tested, apart from CoA. Morphologically, the M2 subpopulation represents the "classical" type 1 mitral cells, the large neurons whose somata and lateral dendrites lie deep in EPL (Mori *et al*., 1983; Orona *et al*., 1984). The M2 subtype therefore corresponds to the population with the most widespread projections (Chen *et al*., 2022; Igarashi *et al*., 2012). This subtype may therefore serve to broadcast the extracted information throughout the cortex. Consistent with previous studies, the projection pattern for CoA suggests that the pathway involving CoA is specialized (Chen *et al*., 2022; Sosulski *et al*, 2011).

As expected from projection bias to anterior areas (Chen *et al*., 2022; Haberly & Price, 1977; Igarashi *et al*., 2012), tufted cell subtypes tended to be labelled by retrograde labeling from more anterior coordinates. In general, restricting the retrograde labeling to Cck-expressing neurons revealed morphologically heterogeneous subsets of tufted cells.

Neurons expressing ET2 markers were selective for AONpe. Populations most enriched in the T1 markers differed from ET subtypes, with the T1 marker transcripts also enriched in OTcap-projecting neurons. This population therefore may correspond to the OT-projecting tufted cells identified by the MAPseq method (Chen *et al*., 2022). Markers for the ET4 subtype were not robustly enriched in our samples. Given that the ET4 markers are enriched in the Cck-RIP samples of RiboTag mice, other injection coordinates in Cck-Cre may reveal this subtype.

One limitation of our study is that the description was at the population level. The average description of transcriptomic data may cause minor populations to go undetected. For example, in our Cck-Cre::Ribotag mouse data, the ET2 markers were undetected. ET2 marker transcripts could be detected only when they were enriched using the retrograde labeling method. Previous works that used high-throughput methods, such as single-nucleus sequencing or projection-based sequencing methods provide a wealth of resources for the sparser populations that have not been resolved here.

Physiologically, the large mitral cells labelled by the Tbx21-aPIR injections deviated most from the population average. We further note that they have the weakest baseline sniff locking. The surprising uniformity of preferred phase in the general population suggests that extracellular neurophysiology may over-represent tufted cells. We speculate that this may explain why it has been difficult to observe baseline sniff-phase differences using extracellular recording methods. The ability to optogenetically tag a subpopulation therefore gives the field an opportunity to study previously under-represented cell types to be characterized. It will be an intriguing future study to investigate how specific subtypes of olfactory bulb outputs contribute to olfactory processing, especially in behaving animals, which may be subject to differential modulatory influences (Chae *et al*., 2022; Kapoor *et al*., 2016; Otazu *et al*., 2015; Yamada *et al*., 2017).

Overall, our findings indicate that downstream olfactory regions do not receive a uniform output from a single olfactory bulb neuron class. Instead, overlapping subsets are innervated by genetically accessed projection neuron populations with distinct transcriptomic signatures, anatomical organization, and intrinsic physiological properties. This combinatorial diversity reveals an additional level of organization in the second stage of olfactory processing and may be a mechanism through which olfactory features are differentially routed and transformed into percepts and behavioural outcomes.

## Acknowledgement

We thank Yu-Pei Huang for technical support and Sayori Gordon for administrative contributions. We are grateful to Marco Terenzio for helpful suggestions on the RiboTag method and Xiang-Chun Ju and all members of the lab for comments on the project. We are grateful for the help and support provided by the staff of Animal Resources Section, Scientific Imaging section, Sequencing section, Scientific Computing and Data Analysis section, and Basic Lab Support section of Core Facilities at OIST Graduate University. This work was supported by JSPS KAKENHI Grant Number JP24K09688 and OIST Graduate University.

## Methods

### Animals

All experiments were conducted in AAALAC-approved facility and all protocols had been approved by OIST’s Animal Care and Use Committee (Protocol number: ACUP-2026-054). Tbx21-Cre (B6;CBA-Tg(Tbx21-cre)1Dlc/J; JAX #024507, (Haddad *et al*., 2013)), Lbhd2-CreERT2 (C57BL/6J-Lbhd2^em1(cre/ERT2)Fuki^/J; JAX #036054, (Koldaeva *et al*., 2021)) and CCK-IRES-Cre (Ccktm^1.1(cre)Zjh^/J; JAX #012706, (Taniguchi *et al*., 2011)). For the crossed mice, these Cre mice were mated with Ai14 reporter mice (B6.Cg-Gt(ROSA)26Sor^tm14(CAG-tdTomato)HZE^/J;JAX #007914, (Madisen *et al*, 2010)) or RiboTag mice (B6N.129-Rpl22^tm1.1Psam/J^; JAX #011029, (Sanz *et al*, 2019)). Both male and female mice in Bl6C57/J background were used for the experiments. Virus injections were performed on mice at 8-10 weeks age. All experimental and control mice were perfused within the 8-14 weeks age range. Lbhd2-CreERT2::Ai14 and Lbhd2-CreERT2::RiboTag mice were administered Tamoxifen via food at P21 according to the protocol described previously (Fu *et al*, 2023).

### Virus constructs

Recombinant retrograde AAVs (AAVrg (Tervo *et al*, 2016)) were synthetised by Vector Builder (Kanagawa, Japan). The following AAVrg (serotype 2) were used: AAVrg-CAG-GFP (Addgene: 37825-AAVrg; lot v145251; titer: 1.9 x 10^13^ GC/ml), AAVrg-FLEX-RPL22HA-eYFP (Addgene, titer 1.11 x 10^14^, lot# 240706AAVD09, titre 1.26 x 10^14^, lot# 241025AAVP11), AAVrg-EF1a-FLEX-ChR2-eYFP (Addgene, 1.0e14 GC/ml, lot# 250506AAVN15). Upon delivery, the viruses were aliquoted and stored at -80°C for long term storage. The 5 µl working aliquots were stored at 4°C.

### Stereotaxic viral injections

Virus was backfilled into a glass capillary (3.5" Drummond #3-000-203-G) pre-filled with mineral oil (M5904, Sigma-Aldrich/Merck, Germany) and pulled with a vertical puller (PC-100, Narishige, Japan). Mice were anesthetized using isoflurane (3% induction, 1.5% maintenance; IsoFlo, Zoetis Japan) and transferred to a stereotaxic frame (Model 1900, Kopf Instruments, USA). Body temperature was maintained throughout surgery at approximately 37°C via heating pad and thermostat (#40-90-8D, FHC USA). The fur covering the head was shaved, and the exposed scalp sterilized using 10% povidone-iodine solution (Shionogi, Japan) and incised to expose the frontal bone. The Lambda and Bregma landmarks were levelled, and the target stereotactic coordinates were marked on the skull surface (see table 4 for the stereotaxic coordinates). Craniotomy was made using a dental drill (Osada Success 40). Prior to virus injection, the glass capillary was coated with DiI (V222885, Invitrogen/ThermoFisher Scientific, USA) to 2 mm above the tip. After lowering the capillary tip to the coordinate, the virus was dispensed using a Nanoject III (Drummond Scientific, USA) unilaterally for AAVrg-CAG-GFP and AAVrg-EF1a-FLEX-ChR2-eYFP, and bilaterally for AAVrg-EF1a-FLEX-Rpl22HA-eYFP. The volume of the virus was adjusted according to the size of the target structure and presence of other adjacent olfactory areas. The capillary was left in place for 5 minutes after the injection before retraction. The craniotomy was sealed with silicone sealant (Kwik-Cast, World Precision Instruments) before suturing the scalp. Animals were kept in a warm recovery chamber post-operatively for approximately 1.5 hours and then returned to the cage. Carprofen (Rimadyl, Zoetis, Japan) was administered intraperitonially (5mg/kg) during the surgery and post-operatively for 3 days.

**Table 4.** Injection conditions used in the study.

| Virus and injection type | Cortical target | Injection co-ordinates (ML, AP, DV) in mm rel. Bregma | Volume injected (nL) | Speed of injection | Incubation time in weeks |
|---|---|---|---|---|---|
| AAVrg-CAG-GFP (unilateral) | AON | 1.2, 3.0, 3.3 | 24 | 0.2nl/sec | 3 |
|  | OT | 1.4, 1.4, 5.65 | 100 | 0.4nl/sec | 4 |
|  | PIR | 3.8, -1.5, 4.8 | 100 | 0.4nl/sec | 4 |
| AAVrg-FLEX-RPL22HA-eYFP (bilateral) | AONpe | ±1.35, 2.5, 3.2 (at 15º angle wrt DV axis) | 24 | 0.2nl/sec | 3 |
|  | AONvl | ±1.2, 2.7, 4.1 | 24 | 0.2nl/sec | 3 |
|  | OTcap | ±0.76, 2.2, 5.0 | 100 | 0.4nl/sec | 3 |
|  | aPIR | ±3.0, 0.3, 5.38 | 24 | 0.2nl/sec | 3 |
|  | pPIR | ±4.0, -1.5, 4.8 | 100 | 0.4nl/sec | 4 |
|  | CoA | ±2.8, -1.8, 5.75 | 100 | 0.4nl/sec | 4 |
| AAVrg-EF1a-FLEX-ChR2-eYFP (unilateral) | AONpe | 1.35, 2.5, 3.2 (at 15º angle wrt DV axis) | 24 | 0.2nl/sec | 2 |
|  | OTcap | 0.76, 2.2, 5.0 | 100 | 0.4nl/sec | 3 |
|  | aPIR | 3.0, 0.3, 5.38 | 24 | 0.2nl/sec | 3 |

### Histology

2-4 weeks after virus injection (Table 4), mice were deeply anesthetized with isoflurane until cessation of breathing and perfused transcardially with 0.1 M phosphate buffer (PB; 22.6 mM NaH_2_PO_4_, 77.4 mM Na_2_HPO_4_, pH7.4; 05071, Yoneyama Chemical and 28-3760-5, Sigma-Aldrich/Merck, respectively) followed by 4% paraformaldehyde (PFA) solution in PB. Brains were removed from the skull and post-fixed in 4% PFA overnight at 4 °C, and washed in PB before sectioning. Each brain was aligned using a custom mold, cut sagittally into two hemispheres, and embedded in 4% agarose (Sigma-Aldrich, A9539). Coronal sections (100 µm) were cut on a vibratome (5100mz-Plus, Campden Instruments, UK) and collected in PB. For the OB, every fourth section was collected (six sections per hemisphere) to sample the anteroposterior extent of the OB. For sections containing the injection sites, all sections containing the DiI signal were collected. Selected sections were mounted onto coated glass slides (Matsunami Glass, 3-8437-01).

### Immunohistochemistry

Sections were submerged for 3 hours at room temperature in blocking buffer (5% goat serum from Sigma-Aldrich, G6767, in PB containing 0.1% Triton X-100 from Sigma-Aldrich, X100). They were then incubated overnight at 4 °C with chicken anti-GFP primary antibody (1:500 in blocking buffer; Abcam, ab13970). The following day, sections were washed with PBT (3 x 5 min) and incubated for 1h at RT with Alexa Fluor 488-conjugated goat anti-chicken secondary antibody (1:300 in blocking buffer; Abcam, ab150169). After a 15-minute wash in PBT, nuclei were counterstained with DAPI (1:500 in PB; Sigma-Aldrich, D9542) for 5 minutes. Slides were mounted with Fluoromount-G (Invitrogen, 00-4958-02).

### Confocal imaging

Images were acquired on an LSM 900 confocal microscope (Carl Zeiss, Germany) using the 405 nm (for DAPI), 488 nm (for GFP/EYFP) and 561 nm (for tdTomato and DiI) lasers. Stained sections were scanned with a 10x objective (Plan-APOCHROMAT 10x/0.45, Zeiss, Germany) for overview of the injection sites or with a 20x objective (Plan-APOCHROMAT 20x/0.17, Zeiss) for morphological analysis of fluorescence labelling patterns in the olfactory bulb. Coronal sections containing the injection sites were additionally scanned around the area containing the LOT using the 20x objective. Resolution used for 20x scans was 1024 x 1024 pixels (1 pixel = 0.312µm x 0.312 µm) and for 10x was 768 x 768 pixels (1 pixel = 0.832 µm x 0.832 µm) and overlaid with a stitching function in the associated software (ZEN blue) to obtain a tiled image.

### RNA-immunoprecipitation (RNA-IP)

Mice were transcardially perfused with ice-cold PB, then the OBs were rapidly dissected, snap-frozen on dry ice, then stored at -80°C. For mice injected with AAVrg, the rest of the brain was incubated overnight at 4°C in 4% PFA in PB, then sectioned and imaged to confirm expression of eYFP at the correct location.

RNA-IP was performed according to established protocols (Sanz et al 2015; Sanz et al 2019). Buffers for RNA-IP were freshly made on the day of use and stored on ice. Tissue was homogenized in 500 µl ice-cold homogenization buffer (50 mM Tris, pH 7.5, 100 mM KCl, 12 mM MgCl_2_, 1% NP-40, 1 mM DTT, 200 U/ml RNaseOUT, 1 mg/ml heparin, 100 µg/ml cycloheximide, plus protease inhibitors; Sanz et al, 2019) by passing it through a 20-gauge needle 10 times. Homogenates were mixed gently for 30 minutes at 4°C, and then centrifuged at 10,000 G for 10 minutes at 4°C. 400 µl of the cleared lysate was transferred to a clean RNase-free tube. A 20 µl (5%) input sample was removed into a second tube, frozen on dry ice and stored at -80°C. 3 µl of mouse anti-HA antibody (#901513; BioLegend) was added to each the remaining 380 µl of cleared lysates and samples were mixed for 4 hours at 4°C. The lysates containing the antibody were transferred to tubes containing 150 µl of washed Protein A/G-conjugated magnetic beads (#88803; Pierce) and mixed gently overnight at 4°C.

Tubes were placed in a magnetic stand for 1 minute and the supernatant removed. The beads were washed four times in 800 µl freshly prepared high-salt wash buffer (50 mM Tris, pH 7.5, 300 mM KCl, 12 mM MgCl_2_, 1% NP-40, 0.5 mM DTT, 100 µg/ml cycloheximide) for 5 minutes each at 4°C with gentle mixing. After the final wash, the beads and wash buffer were transferred to a clean tube to minimize carryover of non-specific RNA bound to the wall of the tube. The wash buffer was removed, and 300 µl RNA lysis buffer (Monarch Total RNA Miniprep Kit; T2010S, New England Biolabs) was added. The samples were vortexed for 20 seconds at room temperature, then placed in the magnetic stand and the buffer transferred to a clean tube. 300 µl RNA lysis buffer was also added to the 20 µl input samples taken the previous day and samples were vortexed.

RNA from all input and IP samples was isolated using a Monarch Total RNA Miniprep Kit following the manufacturer’s protocol. RNA concentration and quality was assessed using the Agilent 2100 Bioanalyzer and Agilent RNA 6000 Pico kit. Only RNA with an RNA integrity number (RIN) ≥ 8.0 was used for RNA sequencing. Extracted RNA was stored at -80°C before use in RNA sequencing.

### RNA library preparation and sequencing

Poly(A) mRNA was isolated using the NEBNext Poly(A) mRNA Magnetic Isolation Module (New England Biolabs, E7490). Stranded libraries were prepared with the NEBNext Ultra II Directional RNA Library Prep Kit for Illumina (New England Biolabs, E7760) and indexed with NEBNext Multiplex Oligos for Illumina (Unique Dual Index UMI Adaptors RNA Set 1; New England Biolabs, E7416S).

Libraries were sequenced on an Illumina NovaSeq X using a 1.5B flow cell with a 300-cycle reagent kit.

Base call (BCL) files were converted to FASTQ and demultiplexed using Illumina BCL Convert (v4.3.6 and v4.3.13). UMI adapter sequences were added to the header of each read using UMI-tools v1.1.6 (Smith et al, 2017). Read quality filtering and adapter trimming were performed using Trimmomatic v0.39 (Bolger et al 2014) in paired-end mode using the ILLUMINACLIP step with standard TruSeq2 adapters (2:30:10 seed mismatches, palindrome threshold and simple threshold). Low-quality leading and trailing bases below a Phred quality score of 3 were removed (LEADING:3, TRAILING:3). Reads were scanned using a 4-base sliding window and trimmed when the average quality dropped below 15 (SLIDINGWINDOW:4:15), and the first 11bp were removed from the 5’ end of each read (HEADCROP:11). Finally, any resulting reads shorter than 25 bp were discarded (MINLEN:25).

Filtered reads were aligned to the indexed mouse genome (mouse genome assembly GRCm39) using STAR v2.7.7a (Dobin *et al*, 2013). Aligned reads were sorted and indexed using SAMtools v1.12 (Li *et al*, 2009). Deduplication of indexed reads was performed in UMI-tools, before counting with featureCounts in Subread v2.0.2 (Liao *et al*, 2014).

### Acute electrophysiology with opto-tagging

AAVrg-EF1a-FLEX-ChR2-eYFP was injected into the left AONpe or OTcap of Cck-Cre mice or aPIR of Tbx21-Cre mice as described above, and acute electrophysiology was conducted after incubation period.

Optogenetic stimuli were generated with a 450 nm blue laser diode (L450P1600MM, Thorlabs) driven by an LED driver (DC4100, Thorlabs, USA) using a custom software written in LabVIEW (National Instruments, USA) and coupled into an optrode with an optical fiber (0.22 NA, 50/62.5 μm) positioned 0.1 mm above the highest contact site of a 32-channel probe (A1x32-Poly3-5mm-25s-177-OCM32LP, NeuroNexus, USA). The average powers for the 5 ms pulse were 0.012, 0.22, and 0.34 mW, measured by a power meter (PM320E with Silicon power head, S120C, Thorlabs, USA). Extracellular recording and copy of the commands sent to the LED driver were acquired at 30 kHz with an Intan acquisition system (RHD 32ch headstage and RHD512-channel recording controller, Intan Technologies, USA). Respiratory signal and another copy of the LED command were acquired with a data acquisition interface (Power 1401, Cambridge Electronic Design Ltd, UK) with its custom environment (Spike 2, CED). The two datasets were synchronized using the LED command signals.

The mice were anesthetized with isoflurane and placed in a stereotaxic frame. The skin over the frontal bone was excised and the exposed skull was cleaned and dried before a small craniotomy was made to insert a reference pin into the right hippocampal area and attach an aluminum headplate with gel superglue and dental acrylic. A small craniotomy (∼ 0.5 mm) was made over the dorsolateral olfactory bulb, targeting the medio-lateral level where the mitral cell layer runs vertically. Care was taken to avoid surface damage to preserve the integrity of the superficial layer, which was confirmed by minimal or no bleeding and intact blood vessels on the surface. The animal was removed from isoflurane, administered with ketamine/xylazine intraperitoneally (ketamine, 100 mg/kg and xylazine, 20mg/kg) and transferred to a recording set up. The body temperature was maintained at 37°C. Respiration was monitored continuously by placing a flow sensor (AWM3100V, Honeywell, USA) next to the right nostril. The extracellular recording probe was lowered to 0.3 mm below the surface using a micromanipulator (3-axis Mini25, Luigs & Neumann, Germany). To minimize drift, the recording did not commence until at least 5 minutes after lowering the electrode to a new recording depth. After at least 1 minute of baseline recording, blue light pulses were delivered every 5 seconds with the 3 light levels intermixed and repeated at least 8 times for each light level. Recordings were obtained from multiple depths using the step function (4 μm step). After 1.5 hours, mice were deeply anaesthetised with isoflurane until breathing stopped and perfused transcardially with 0.1 M PB and 4 % PFA. The head was removed and fixed in 4% PFA at 4 °C. After 24 hours, the brain was removed and stored in 0.1 M PB for 24 hours prior to sectioning.

### Analyses

#### Image analysis

Soma segmentation was done on maximum intensity projections of the green and red channels (eYFP and tdTomato fluorescence) of OB images. AI Image Analysis Software (Aivia, Leica Microsystems) was used, with manual training from painting 5-10 somata on example images. Separate training for different injection condition was sometimes necessary due to differences in the density of labelled structures. Detection parameters such as object size range and detection probability threshold were then adjusted to improve the detection accuracy and outline editor tool was used to remove obvious artifacts. The quality of segmentation was checked independently by two people.

EPL boundaries were manually delineated using the DAPI signals only using the polygon selections tool on ImageJ (National Institutes of Health, USA) and the points of the polygons were exported. Using a custom MATLAB (Mathworks, USA) script, the depth of soma relative to the EPL was calculated, where the inner EPL boundary is considered as 0 and the outer EPL boundary is considered as 1. For dendritic signal density, the background was subtracted in ImageJ with the rolling ball radius of 5 pixels, filtered using the "*Tubeness*" function with the sigma parameter set to 0.0033, and finally binarized with a single threshold for all images. The EPL depths were interpolated into 200 bins and the density of binarized signal was recorded.

### Analysis of RNA-seq data

Transcript enrichment in RIP samples relative to their corresponding input samples was analysed using DESeq2 v1.44.0 (Love *et al*, 2014) in R v4.4.3 .and RStudio v2024.12.1. Reads that did not have at least 10 counts in at least 4 samples were excluded from the analysis. The differential expression analysis was followed a shrinkage algorithm to filter noisy data using Approximate Posterior Estimation for Generalized Linear Models (*apeglm* https://github.com/azhu513/apeglm, (Zhu *et al*, 2019)) for the individual injection conditions. For identification of transcripts enriched specifically in CoA RIP samples with respect to the input and other injection conditions, Adaptive SHrinkage with Empricial Bayes (*ashr,* https://github.com/stephens999/ashr (Stephens, 2017)) was applied. Those genes with FDR-adjusted p-values < 0.001 and log2FC > 2 were defined as enriched. Enrichment analysis of subtype-specific marker sets was performed using the Correlation Adjusted MEan RAnk (CAMERA) algorithm (Wu & Smyth, 2012), which analyses the ranks of gene sets in a competitive manner while accounting for inter-gene correlation, with the inter gene correlation set to 0.01 and mean-variance trend set to TRUE. Correction for multiple comparison used the Benjamini-Hochberg procedure.

### Analysis of neurophysiology data

Spike sorting: Spikes were sorted as described previously (Fu *et al*, 2025). Briefly, units were sorted offline using Kilosort 4.0 (Pachitariu et al., 2024), with default parameters from the authors of Kilosort. The outputs from Kilosort were manually curated in the Phy environment (Rossant et al., 2016). Only those clusters with a clear refractory period, with waveforms that were distinguishable from the background, and with stable waveform amplitudes for the majority of the recording were labelled as good single units and used for further analysis.

Classification of opto-tagged units: Opto-tagged units were defined as units exhibiting a tightly locked, short-latency excitatory response to light stimulation. Specifically, a unit was classified as opto-tagged if its peri-stimulus time histogram bin count within a 2 ms - 10 ms window after the onset of the 0.335 mW pulse was more than 2 standard deviations above the baseline.

### Sniff locking analysis

For each sorted unit, recorded spike times were expressed as the phase (in radians) with respect to the sniff cycle. Each cycle was defined from one inhalation peak to the next inhalation peak from the nasal flow record. The preferred phase was the angle of the vector sum, atan2(Σsin (θ*_i_*), Σcos(θ*_i_*)), where θ*_i_* is the phase of individual spikes with respect to the sniff cycle.

## Supplementary figures

**Supplementary Figure 1:**
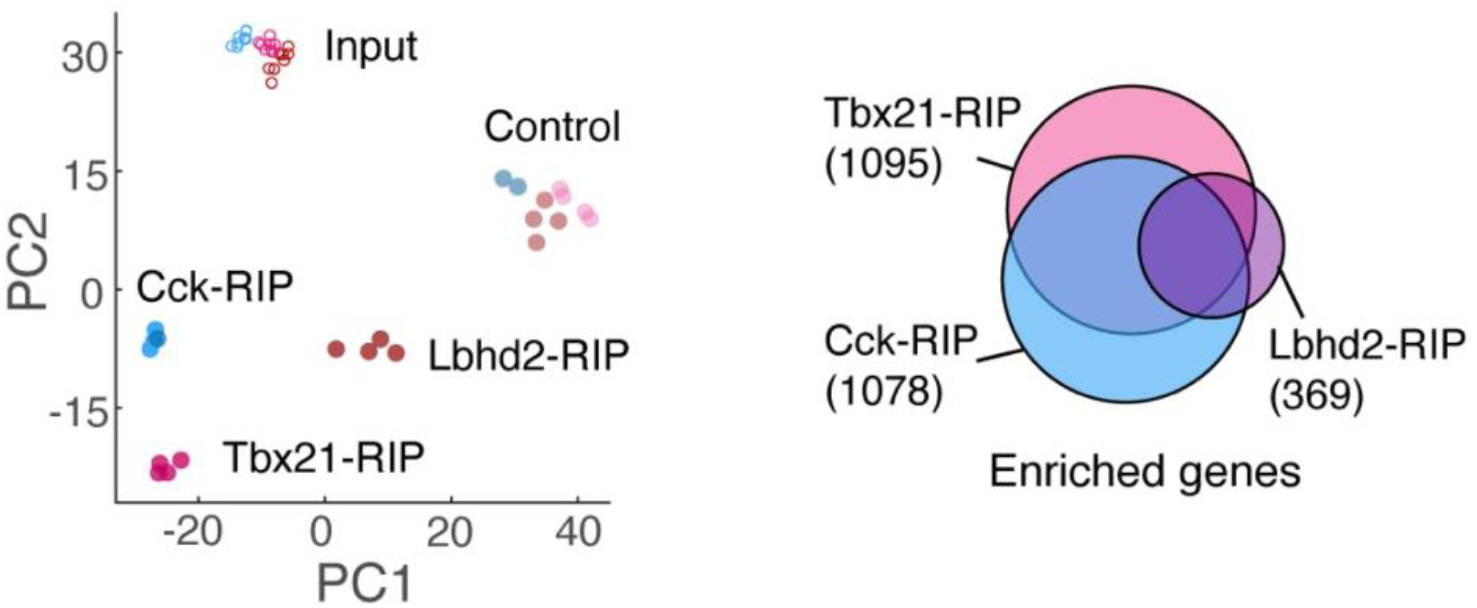
Enrichment of transcripts associated with mitral and tufted cells in Cck-, Tbx21-, and Lbhd2-RIP samples. **A**, Transcript enrichment in RIP samples separates well from those of the input samples, as well as control samples where HA-tagged RPL22 is absent. Principal component axes were calculated from variance-stabilizing transformed count using DESeq2. **B**, Euler plot showing the sets of unique and overlapping enriched transcripts across the three Cre lines. Enriched genes were defined as those with log2FC > 2 (RIP vs. input), and with adjusted p-values less than 0.001.

**Supplementary Fig. 2:**
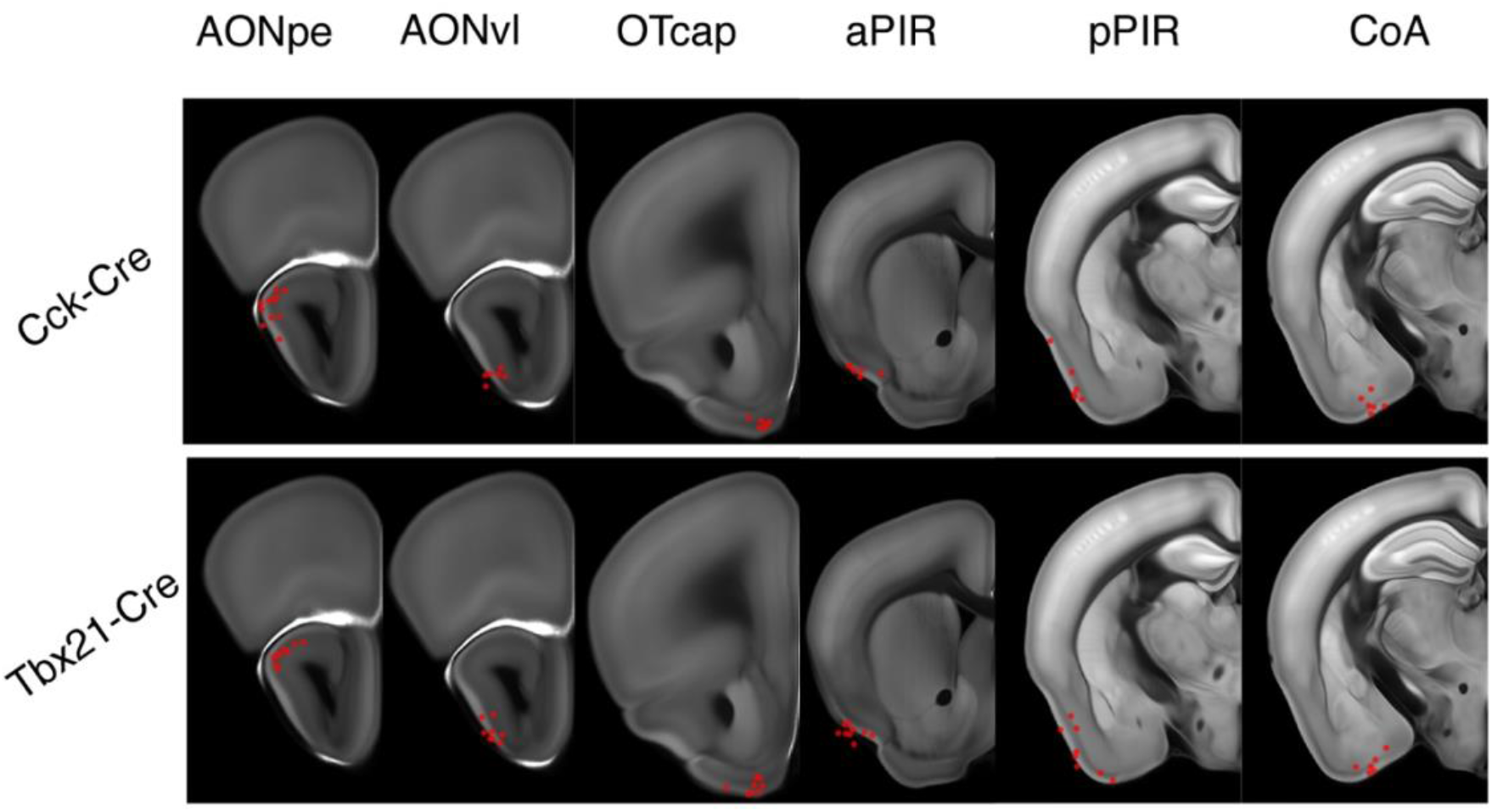
Reconstructed injection coordinates. Locations of estimated injection sites (red dots) on a reference brain co-ordinate (Allen CCF). The tip location of injection coordinate was estimated using a combination of red fluorescence from DiI that coated the injection capillary glass, and the EYFP signal strength especially in the LOT and molecular layer. The injector track signal and DAPI signals were transformed to fit the reference frame using the alignment tool HERBS (Fuglstad *et al*, 2023), and the standardized coordinates were extracted.

**Supplementary figure 3:**
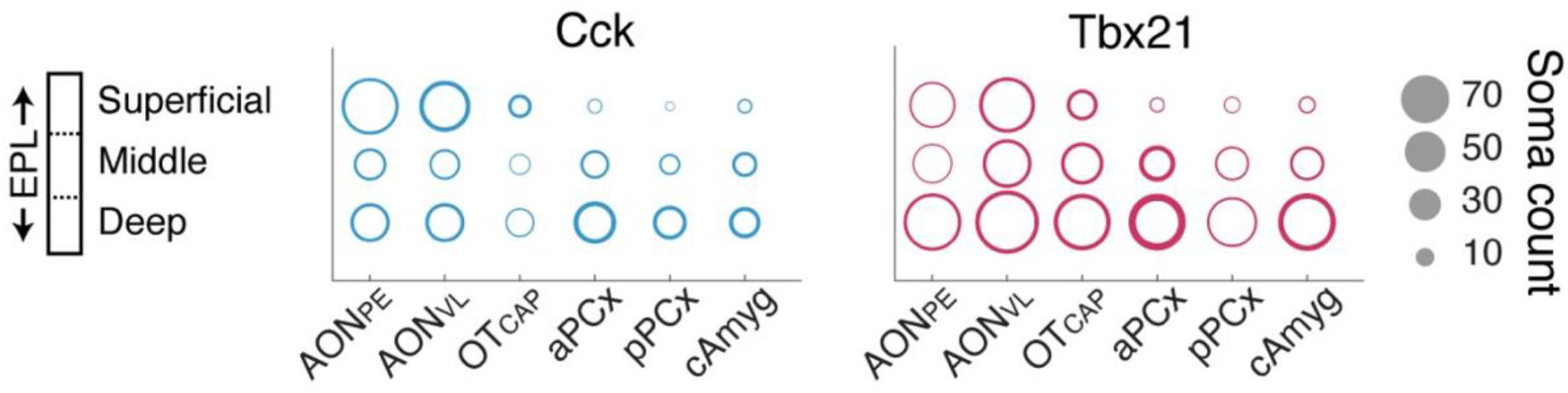
Soma counts sorted by anterior-posterior location of injections & Cre lines. Summary of labelled soma counts for the indicated injection conditions. Somata were segmented using Aivia on fluorescence images acquired by a confocal microscope, and its position expressed as the normalized depth relative to the boundaries of the external plexiform layer. EPL depth was divided into three levels to classify these segmented cells into deep, middle, and superficial somata. The sizes of circles correspond to the numbers of detected somata, with the thickness indicating deviations from the mean (1 standard deviation in each direction, i.e., plus and minus the mean).

**Supplementary figure 4:**
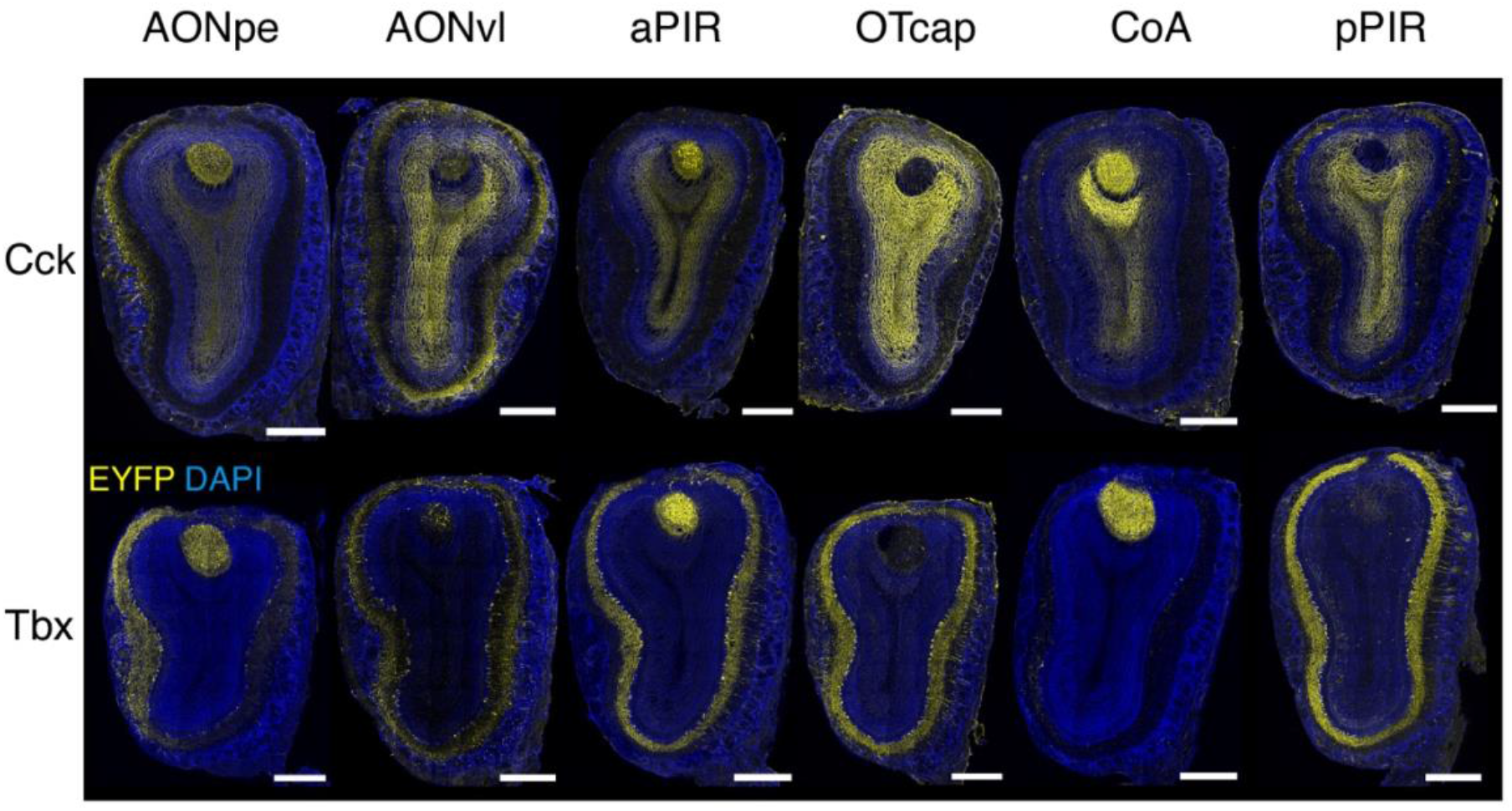
AOB labelled with AONpe, aPIR, and CoA injections. Fluorescent confocal images of coronal sections of olfactory bulb at the level of the AOB (2.4 mm from front), following the injection conditions indicated. Scale bar = 0.5 mm.

**Supplementary figure 5:**
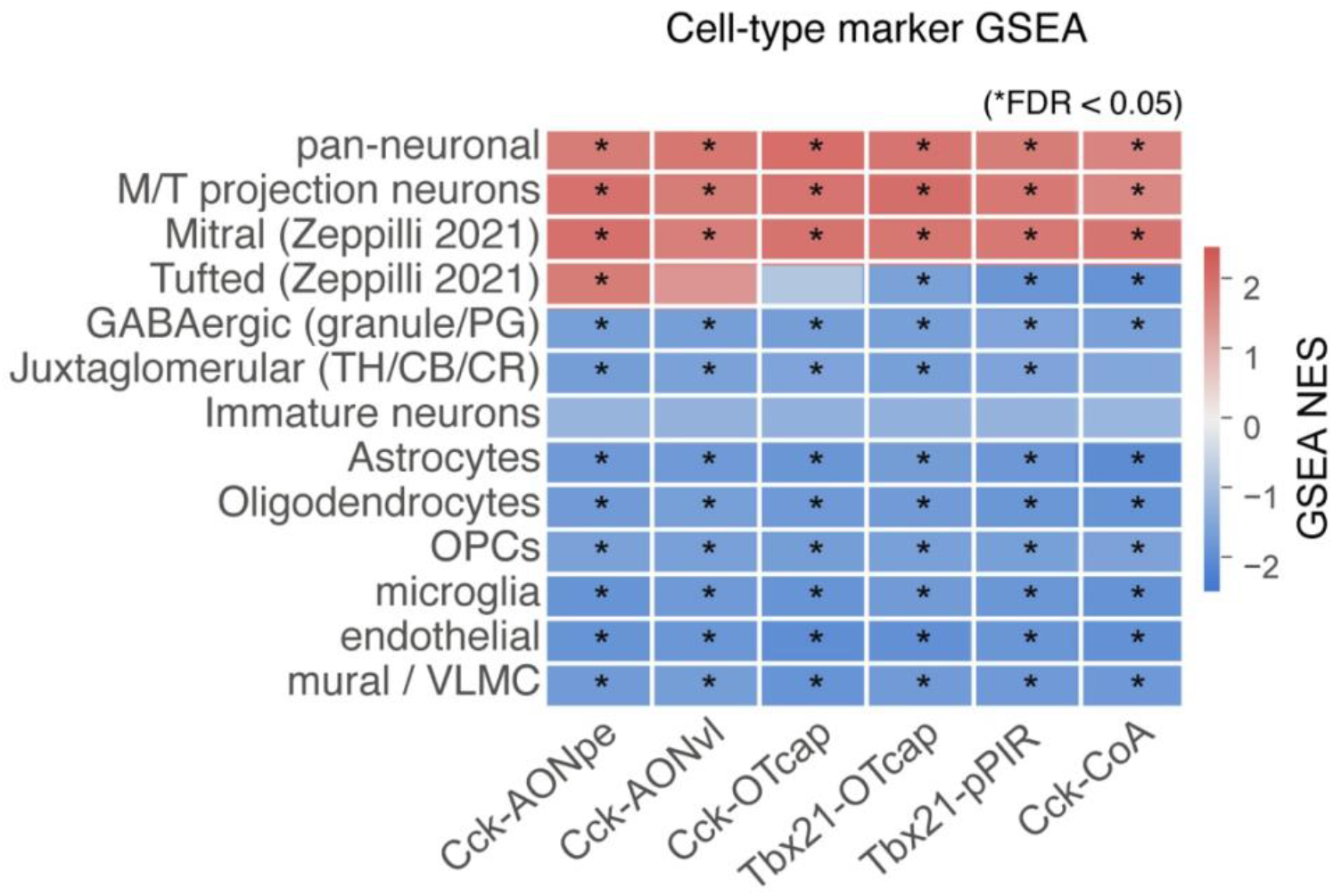
Cell-type marker enrichment confirms projection-neuron-specific IP in injections. Preranked Gean Set Enrichment Analysis (GSEA (Subramanian *et al*, 2005)) of curated olfactory bulb and glial and vascular marker sets (Table 1) in IP versus Input for each condition. Normalized enrichment score (NES) is shown with corresponding colormap, with enrichment and depletion represented by red and blue hues, respectively. The Cre driver genes (Cck, Tbx21) were excluded from all sets.

**Supplementary figure 6:**
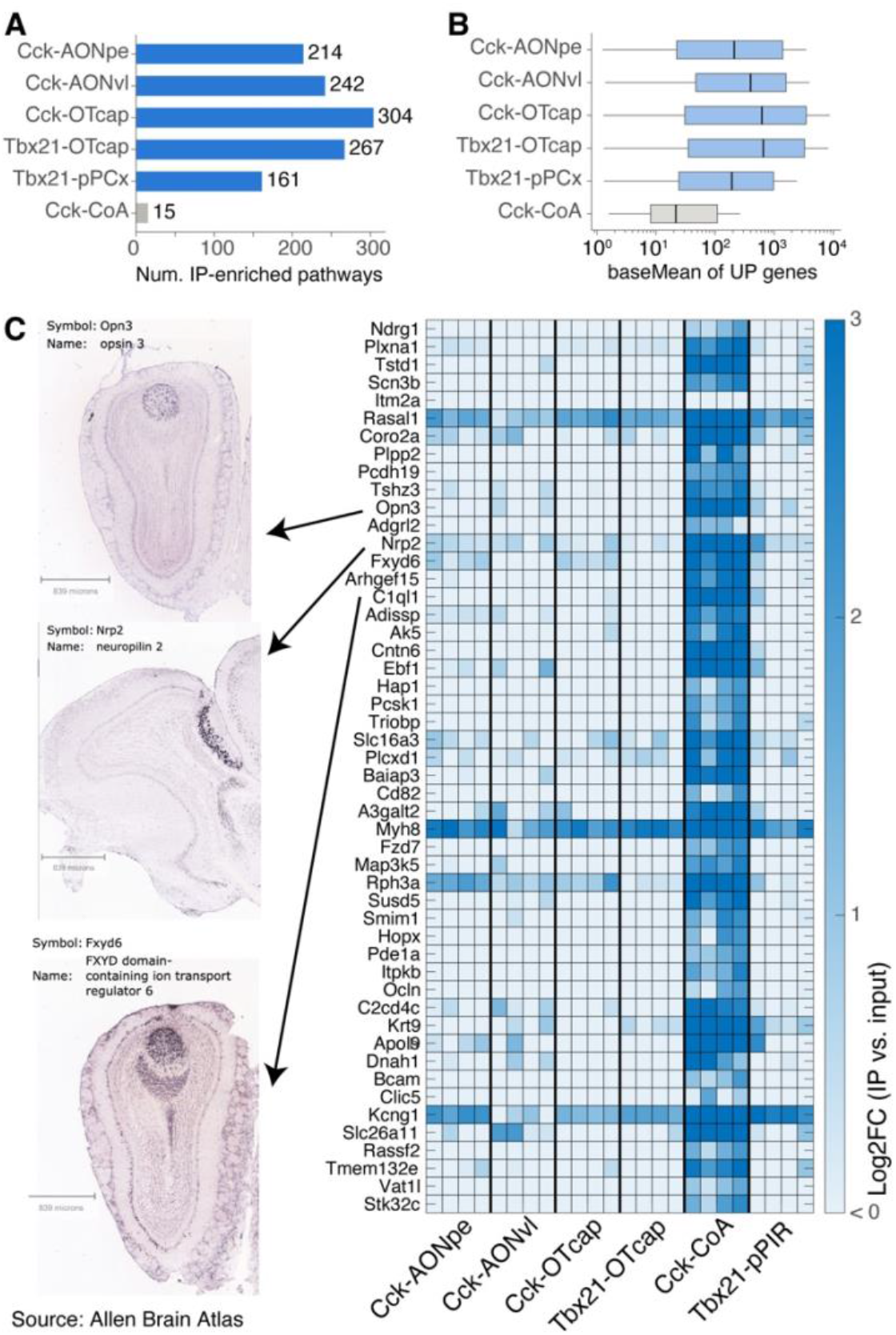
genes enriched in Cck-CoA samples label AOB cells. **A,** Number of gene sets significantly enriched in the RiboTag IP compaired to Input in each condition (preranked GSEA on apeglm-shrunken log2 fold change; NES > 0, BH-adjusted FDR < 0.05). Counts are summed across MSigDB Hallmark, GO:BP/CC/MF, Reactome and KEGG (Legacy and MEDICUS, mapped to mouse orthologs) collections. **B,** Expression level (DESeq2 baseMean, log scale) of IP-enriched genes (padj < 0.05, log2 fold change > 1) in each condition. Boxes show the median and interquartile range, and whiskers extend to 1.5× IQR; outliers are not shown. **C**, Right, Top 50 genes sorted by differential expression in Cck-CoA experiments (relative to all other conditions). See Table 3 for a full list with adjusted p-values, log2 fold change and mean expression levels. Left, In-situ hybridization taken from the Allen Brain Atlas for selected genes (Opn3, Nrp2, and Fxyd6, with original images at https://mouse.brain-map.org/experiment/show/71020433, https://mouse.brain-map.org/experiment/show/80514091, https://mouse.brain-map.org/experiment/show/73592534, respectively).

## Notes

### Competing Interest Statement

The authors have declared no competing interest.

### Summary of Updates

Title, abstract, and one result figure have been edited.

## References

Beiter RM, Rivet-Noor C, Merchak AR, Bai R, Johanson DM, Slogar E, Sol-Church K, Overall CC, Gaultier A (2022) Evidence for oligodendrocyte progenitor cell heterogeneity in the adult mouse brain. Scientific Reports 12: 12921

Chae H, Banerjee A, Dussauze M, Albeanu DF (2022) Long-range functional loops in the mouse olfactory system and their roles in computing odor identity. Neuron 110: 3970–3985.e3977

Chen Y, Chen X, Baserdem B, Zhan H, Li Y, Davis MB, Kebschull JM, Zador AM, Koulakov AA, Albeanu DF (2022) High-throughput sequencing of single neuron projections reveals spatial organization in the olfactory cortex. Cell 185: 4117–4134.e4128

Diamond ME, von Heimendahl M, Knutsen PM, Kleinfeld D, Ahissar E (2008) ’Where’ and ‘what’ in the whisker sensorimotor system. Nature Reviews Neuroscience 9: 601–612

Dobin A, Davis CA, Schlesinger F, Drenkow J, Zaleski C, Jha S, Batut P, Chaisson M, Gingeras TR (2013) STAR: ultrafast universal RNA-seq aligner. Bioinformatics 29: 15–21

Eiting TP, Wachowiak M (2020) Differential Impacts of Repeated Sampling on Odor Representations by Genetically-Defined Mitral and Tufted Cell Subpopulations in the Mouse Olfactory Bulb. The Journal of Neuroscience 40: 6177

Fu X, Huang Y-P, Lindeman S, Mago A, Fukunaga I (2023) Dense and Sparse Labelling of Mitral Cells by Oral and Intraperitoneal Routes of Tamoxifen Administration. In: The Olfactory System, Goldstein B.J., Matsunami H. (eds.) pp. 185-193. Springer US: New York, NY

Fu X, Reinert JK, Reuschenbach J, Mago A, Lindeman S, Fukunaga I (2025) Decoding olfactory bulb output: A behavioral assessment of rate, synchrony, and respiratory phase coding. iScience 28

Fuglstad JG, Saldanha P, Paglia J, Whitlock JR (2023) Histological E-data Registration in rodent Brain Spaces. eLife 12: e83496

Fukunaga I, Berning M, Kollo M, Schmaltz A, Schaefer Andreas T (2012) Two Distinct Channels of Olfactory Bulb Output. Neuron 75: 320–329

Fukunaga I, Herb JT, Kollo M, Boyden ES, Schaefer AT (2014) Independent control of gamma and theta activity by distinct interneuron networks in the olfactory bulb. Nature Neuroscience 17: 1208–1216

Grothe B, Pecka M, McAlpine D (2010) Mechanisms of Sound Localization in Mammals. Physiological Reviews 90: 983–1012

Haberly LB, Price JL (1977) The axonal projection patterns of the mitral and tufted cells of the olfactory bulb in the rat. Brain Research 129: 152–157

Hack MA, Saghatelyan A, de Chevigny A, Pfeifer A, Ashery-Padan R, Lledo P-M, Götz M (2005) Neuronal fate determinants of adult olfactory bulb neurogenesis. Nature Neuroscience 8: 865–872

Haddad R, Lanjuin A, Madisen L, Zeng H, Murthy VN, Uchida N (2013) Olfactory cortical neurons read out a relative time code in the olfactory bulb. Nature Neuroscience 16: 949–957

Igarashi KM, Ieki N, An M, Yamaguchi Y, Nagayama S, Kobayakawa K, Kobayakawa R, Tanifuji M, Sakano H, Chen WR et al (2012) Parallel Mitral and Tufted Cell Pathways Route Distinct Odor Information to Different Targets in the Olfactory Cortex. The Journal of Neuroscience 32: 7970

Kalucka J, de Rooij LPMH, Goveia J, Rohlenova K, Dumas SJ, Meta E, Conchinha NV, Taverna F, Teuwen L-A, Veys K et al (2020) Single-Cell Transcriptome Atlas of Murine Endothelial Cells. Cell 180: 764–779.e720

Kapoor V, Provost AC, Agarwal P, Murthy VN (2016) Activation of raphe nuclei triggers rapid and distinct effects on parallel olfactory bulb output channels. Nature Neuroscience 19: 271–282

Koldaeva A, Zhang C, Huang Y-P, Reinert JK, Mizuno S, Sugiyama F, Takahashi S, Soliman T, Matsunami H, Fukunaga I (2021) Generation and Characterization of a Cell Type-Specific, Inducible Cre-Driver Line to Study Olfactory Processing. The Journal of Neuroscience 41: 6449

Li H, Handsaker B, Wysoker A, Fennell T, Ruan J, Homer N, Marth G, Abecasis G, Durbin R, Genome Project Data Processing S (2009) The Sequence Alignment/Map format and SAMtools. Bioinformatics 25: 2078–2079

Liao Y, Smyth GK, Shi W (2014) featureCounts: an efficient general purpose program for assigning sequence reads to genomic features. Bioinformatics 30: 923–930

Lindeman S, Fu X, Reinert JK, Fukunaga I (2024) Value-related learning in the olfactory bulb occurs through pathway-dependent perisomatic inhibition of mitral cells. PLOS Biology 22: e3002536

Lledo P-M, Alonso M, Grubb MS (2006) Adult neurogenesis and functional plasticity in neuronal circuits. Nature Reviews Neuroscience 7: 179–193

Lois C, Alvarez-Buylla A (1994) Long-Distance Neuronal Migration in the Adult Mammalian Brain. Science 264: 1145–1148

Long JE, Garel S, Alvarez-Dolado M, Yoshikawa K, Osumi N, Alvarez-Buylla A, Rubenstein JLR (2007) Dlx-Dependent and -Independent Regulation of Olfactory Bulb Interneuron Differentiation. The Journal of Neuroscience 27: 3230

Love MI, Huber W, Anders S (2014) Moderated estimation of fold change and dispersion for RNA-seq data with DESeq2. Genome Biology 15: 550

Luskin MB, Price JL (1982) The distribution of axon collaterals from the olfactory bulb and the nucleus of the horizontal limb of the diagonal band to the olfactory cortex, demonstrated by double retrograde labeling techniques. Journal of Comparative Neurology 209: 249–263

Macrides F, Schneider SP (1982) Laminar organization of mitral and tufted cells in the main olfactory bulb of the adult hamster. Journal of Comparative Neurology 208: 419–430

Madisen L, Zwingman TA, Sunkin SM, Oh SW, Zariwala HA, Gu H, Ng LL, Palmiter RD, Hawrylycz MJ, Jones AR et al (2010) A robust and high-throughput Cre reporting and characterization system for the whole mouse brain. Nature Neuroscience 13: 133–140

Marques S, Zeisel A, Codeluppi S, van Bruggen D, Mendanha Falcão A, Xiao L, Li H, Häring M, Hochgerner H, Romanov RA et al (2016) Oligodendrocyte heterogeneity in the mouse juvenile and adult central nervous system. Science 352: 1326–1329

Masuda T, Amann L, Sankowski R, Staszewski O, Lenz M, d’Errico P, Snaidero N, Costa Jordão MJ, Böttcher C, Kierdorf K et al (2020) Novel Hexb-based tools for studying microglia in the CNS. Nature Immunology 21: 802-815

Mori K, Kishi K, Ojima H (1983) Distribution of dendrites of mitral, displaced mitral, tufted, and granule cells in the rabbit olfactory bulb. Journal of Comparative Neurology 219: 339–355

Nagayama S, Homma R, Imamura F (2014) Neuronal organization of olfactory bulb circuits. Frontiers in Neural Circuits Volume 8–2014

Nassi JJ, Callaway EM (2009) Parallel processing strategies of the primate visual system. Nature Reviews Neuroscience 10: 360–372

Orona E, Rainer EC, Scott JW (1984) Dendritic and axonal organization of mitral and tufted cells in the rat olfactory bulb. Journal of Comparative Neurology 226: 346–356

Otazu Gonzalo H, Chae H, Davis Martin B, Albeanu Dinu F (2015) Cortical Feedback Decorrelates Olfactory Bulb Output in Awake Mice. Neuron 86: 1461–1477

Parrish-Aungst S, Shipley MT, Erdelyi F, Szabo G, Puche AC (2007) Quantitative analysis of neuronal diversity in the mouse olfactory bulb. Journal of Comparative Neurology 501: 825–836

Pastrana E, Cheng L-C, Doetsch F (2009) Simultaneous prospective purification of adult subventricular zone neural stem cells and their progeny. Proceedings of the National Academy of Sciences 106: 6387–6392

Roska B, Meister M (2014) The retina dissects the visual scene into distinct features. In: The New Visual Neurosciences, Werner J.S., Chalupa L.M. (eds.) pp. 163-182. MIT Press:

Sanes JR, Masland RH (2015) The types of retinal ganglion cells: current status and implications for neuronal classification. Annu Rev Neurosci 38: 221–246

Sanz E, Bean JC, Carey DP, Quintana A, McKnight GS (2019) RiboTag: Ribosomal Tagging Strategy to Analyze Cell-Type-Specific mRNA Expression In Vivo. Current Protocols in Neuroscience 88: e77

Sanz E, Yang L, Su T, Morris DR, McKnight GS, Amieux PS (2009) Cell-type-specific isolation of ribosome-associated mRNA from complex tissues. Proceedings of the National Academy of Sciences 106: 13939–13944

Scott JW (1981) Electrophysiological identification of mitral and tufted cells and distributions of their axons in olfactory system of the rat. Journal of Neurophysiology 46: 918–931

Sosulski DL, Bloom ML, Cutforth T, Axel R, Datta SR (2011) Distinct representations of olfactory information in different cortical centres. Nature 472: 213–216

Stephens M (2017) False discovery rates: a new deal. Biostatistics 18: 275–294

Subramanian A, Tamayo P, Mootha VK, Mukherjee S, Ebert BL, Gillette MA, Paulovich A, Pomeroy SL, Golub TR, Lander ES et al (2005) Gene set enrichment analysis: A knowledge-based approach for interpreting genome-wide expression profiles. Proceedings of the National Academy of Sciences 102: 15545–15550

Taniguchi H, He M, Wu P, Kim S, Paik R, Sugino K, Kvitsani D, Fu Y, Lu J, Lin Y et al (2011) A Resource of Cre Driver Lines for Genetic Targeting of GABAergic Neurons in Cerebral Cortex. Neuron 71: 995–1013

Tasic B, Yao Z, Graybuck LT, Smith KA, Nguyen TN, Bertagnolli D, Goldy J, Garren E, Economo MN, Viswanathan S et al (2018) Shared and distinct transcriptomic cell types across neocortical areas. Nature 563: 72–78

Tepe B, Hill MC, Pekarek BT, Hunt PJ, Martin TJ, Martin JF, Arenkiel BR (2018) Single-Cell RNA-Seq of Mouse Olfactory Bulb Reveals Cellular Heterogeneity and Activity-Dependent Molecular Census of Adult-Born Neurons. Cell Reports 25: 2689–2703.e2683

Tervo DGowanlock R, Hwang B-Y, Viswanathan S, Gaj T, Lavzin M, Ritola Kimberly D, Lindo S, Michael S, Kuleshova E, Ojala D et al (2016) A Designer AAV Variant Permits Efficient Retrograde Access to Projection Neurons. Neuron 92: 372–382

Vanlandewijck M, He L, Mäe MA, Andrae J, Ando K, Del Gaudio F, Nahar K, Lebouvier T, Laviña B, Gouveia L et al (2018) A molecular atlas of cell types and zonation in the brain vasculature. Nature 554: 475–480

Waclaw RR, Allen ZJ, Bell SM, Erdélyi F, Szabó G, Potter SS, Campbell K (2006) The Zinc Finger Transcription Factor Sp8 Regulates the Generation and Diversity of Olfactory Bulb Interneurons. Neuron 49: 503–516

Wassle H (2004) Parallel processing in the mammalian retina. Nat Rev Neurosci 5: 747–757

Wässle H (2004) Parallel processing in the mammalian retina. Nature Reviews Neuroscience 5: 747-757

Wu D, Smyth GK (2012) Camera: a competitive gene set test accounting for inter-gene correlation. Nucleic Acids Research 40: e133–e133

Yamada Y, Bhaukaurally K, Madarász TJ, Pouget A, Rodriguez I, Carleton A (2017) Context- and Output Layer-Dependent Long-Term Ensemble Plasticity in a Sensory Circuit. Neuron 93: 1198–1212.e1195

Yao Z, van Velthoven CTJ, Kunst M, Zhang M, McMillen D, Lee C, Jung W, Goldy J, Abdelhak A, Aitken M et al (2023) A high-resolution transcriptomic and spatial atlas of cell types in the whole mouse brain. Nature 624: 317–332

Zeisel A, Hochgerner H, Lönnerberg P, Johnsson A, Memic F, van der Zwan J, Häring M, Braun E, Borm LE, La Manno G et al (2018) Molecular Architecture of the Mouse Nervous System. Cell 174: 999–1014.e1022

Zeppilli S, Ackels T, Attey R, Klimpert N, Ritola KD, Boeing S, Crombach A, Schaefer AT, Fleischmann A (2021) Molecular characterization of projection neuron subtypes in the mouse olfactory bulb. eLife 10: e65445

Zhu A, Ibrahim JG, Love MI (2019) Heavy-tailed prior distributions for sequence count data: removing the noise and preserving large differences. Bioinformatics 35: 2084–2092

